# MBD8 is required for LDL2-mediated transcriptional repression downstream of H3K9me2 in *Arabidopsis*

**DOI:** 10.1101/2025.08.21.671526

**Authors:** Shusei Mori, Akihisa Osakabe, Juliarni, Yuriko Tanaka, Mami Hirayama, Soichi Inagaki, Tetsuji Kakutani

## Abstract

Histone H3 lysine 9 methylation (H3K9me) is a conserved epigenetic mark for transcriptional gene repression. In Arabidopsis, H3K9 dimethylation (H3K9me2) promotes the removal of a transcription-associated mark, H3K4 monomethylation (H3K4me1), and this process within genes requires the histone demethylase LDL2. However, how LDL2 specifically functions in genes that accumulate H3K9me2 remains unknown. Here, we show that methyl-CpG-binding domain protein 8 (MBD8) binds to LDL2 and plays a role in H3K4me1 removal and transcriptional gene repression downstream of H3K9me2. MBD8 preferentially binds GC-poor DNA, regardless of its cytosine methylation status. Interestingly, LDL2 protein levels were significantly reduced in the *mbd8* mutant background, suggesting that MBD8 stabilizes the LDL2 protein, facilitating H3K4me1 removal and the transcriptional repression of H3K9me-marked genes. The domain necessary for the MBD8-LDL2 interaction is conserved among plants. We propose that MBD proteins have diverse functions beyond their methylated DNA-binding capabilities.

## Introduction

In eukaryotes, chromatin modifications such as DNA cytosine methylation (mC) and histone modifications regulate transcription. The methylations of histone H3 at lysine 4 (H3K4) and lysine 9 (H3K9) are highly conserved chromatin modifications associated with euchromatin and heterochromatin, respectively ^1,2^. H3K9me is involved in the transcriptional silencing of genes and transposons ^3,4^, but how H3K9me represses transcription and interacts with other chromatin modifications remains unclear.

H3K9me has been reported to recruit reader proteins to repress transcription ^5^. In fission yeast and animals, heterochromatin protein 1 (HP1) family proteins recognize H3K9me through their chromodomains and play critical roles in heterochromatin formation and gene silencing ^3,6–8^. In *A. thaliana*, H3K9 dimethylation (H3K9me2) is the major form of H3K9 in constitutive heterochromatin ^5^, and recent screens for H3K9me2 reader proteins identified AGDP1 (also known as ADCP1) ^9,10^. AGDP1 binds to H3K9me2 by its tandem agenet domains (AGDs). Interestingly, H3K4 methylation decreases the binding of AGDP1 to H3K9me2 peptides, suggesting that AGDP1 recognizes H3K9me2 and unmethylated K4 on the H3 tail (H3K4me0). In the *agdp1* mutant, the H3K9me2 level decreases, and transposable elements (TEs) are derepressed. AGDP1 is also required for heterochromatic chromocenter formation ^10,11^. In addition, H3K9me2 recruits the DNA methyltransferases CMT2 and CMT3, which mediate cytosine methylation in the non-CpG context (referred to as mCH, where H can be A, T, or C). CMT2 and CMT3 contain a bromo adjacent homology (BAH) domain and a chromodomain, which cooperatively recognize the H3 tail containing H3K9me2 ^12,13^. In turn, the H3K9 methyltransferases KYP, SUVH5, and SUVH6, harboring the mC-binding SRA domain, are recruited to chromatin with mCH, generating a self-reinforcing loop of H3K9me2 and mCH ^14^. Reader proteins for mC have also been explored, and methyl-CpG-binding domain (MBD) family proteins have been identified ^15,16^. *A. thaliana* has 13 putative MBD protein-encoding genes ^17–19^, although MBD3/13 are pseudogenes ^20–22^. MBD5/6/7 have been confirmed to bind specifically to mC in vitro ^23,24^. MBD1/2/4 do not bind to mC in vitro, but they localize to chromatin with mC in vivo ^25,26^. Specific binding to mC by other MBDs (MBD8/9/10/11) has not been detected ^19,20,23^. Nevertheless, MBD8/9/10/11 have diverse phenotypic effects: MBD8 promotes flowering in the natural accession C24 ^27^; MBD9 is involved in H2A.Z deposition and represses flowering ^28–30;^ MBD10 promotes leaf senescence ^31^; and MBD11 regulates leaf shape and the positioning of flowers ^22^. These observations suggest that these MBDs may contribute to developmental regulation by pathways independent of binding to mC. To elucidate the mechanisms by which H3K9me2 represses transcription, we used the mutant *increase in BONSAI methylation 1* (*ibm1*). The *IBM1* gene encodes a JmjC family histone H3K9 demethylase. In the loss-of-function *ibm1* mutant, H3K9me2 and mCH accumulate ectopically in gene bodies, reflecting the feedback loop between H3K9me2 and mCH ^14,32–34^. The ectopic accumulation of H3K9me2/mCH in *ibm1* leads to transcriptional repression and developmental defects. A previous suppressor screen using developmental defects in *ibm1* as indicators of transcriptional silencing identified the LSD1 family histone demethylase LDL2 ^35^. LDL2 mediates H3K9me2/mCH-induced transcriptional repression by removing monomethylated H3K4 (H3K4me1) in *ibm1*. These results are consistent with our previous studies, which demonstrated tight associations between H3K4me1 and transcription^36,37^. However, the underlying mechanism by which LDL2 specifically functions in H3K9me2/mC-accumulating chromatin to remove H3K4me1 remains unknown.

Here, we identified methyl-CpG-binding protein 8 (MBD8) as a binding partner of LDL2 and revealed that MBD8 is required for the LDL2-induced demethylation of H3K4me1 and associated transcriptional perturbation and developmental defects in the *ibm1* mutant. MBD8 binds to DNA irrespective of cytosine methylation status and is required for LDL2 protein stability. We propose unconventional functions of MBD8 beyond the function as a mC reader.

## Results

### MBD8 binds to LDL2 and mediates the developmental defects in the *ibm1* mutant

To explore LDL2-interacting proteins, we performed immunoprecipitationLJmass spectrometry (IPLJMS) analysis using epitope-tagged LDL2 transgenic lines. The most enriched protein in the immunoprecipitate, except LDL2 itself, was a methyl-CpG-binding domain protein, MBD8 (**Fig. 1a**). To confirm the LDL2-MBD8 interaction, we generated epitope-tagged MBD8 transgenic lines and immunoprecipitated MBD8, and the precipitated proteins were subjected to mass spectrometry. LDL2 was reciprocally detected (**Fig. 1a**), confirming that LDL2 and MBD8 interact with each other in vivo. To examine whether MBD8 functions with LDL2 in the *ibm1* mutant, we crossed *mbd8* T-DNA mutants (*mbd8-3*) with *ibm1-4* ^32^ and generated *ibm1 mbd8-3* double mutants (**Fig. 1b, Extended Data Fig. 1a**). The *ibm1* plants showed developmental defects, such as small, dark green leaves. In contrast, the *ibm1 mbd8* double mutant showed a WT-like phenotype (**Fig. 1b**). Another *mbd8* allele, *mbd8-4*, which we generated via the CRISPR/Cas9 system, also suppressed the developmental defects of *ibm1* (**Extended Data** Fig. 1a, b), confirming that MBD8 functions together with LDL2 to confer developmental defects in the *ibm1* mutant ^35^.

**Figure 1.**
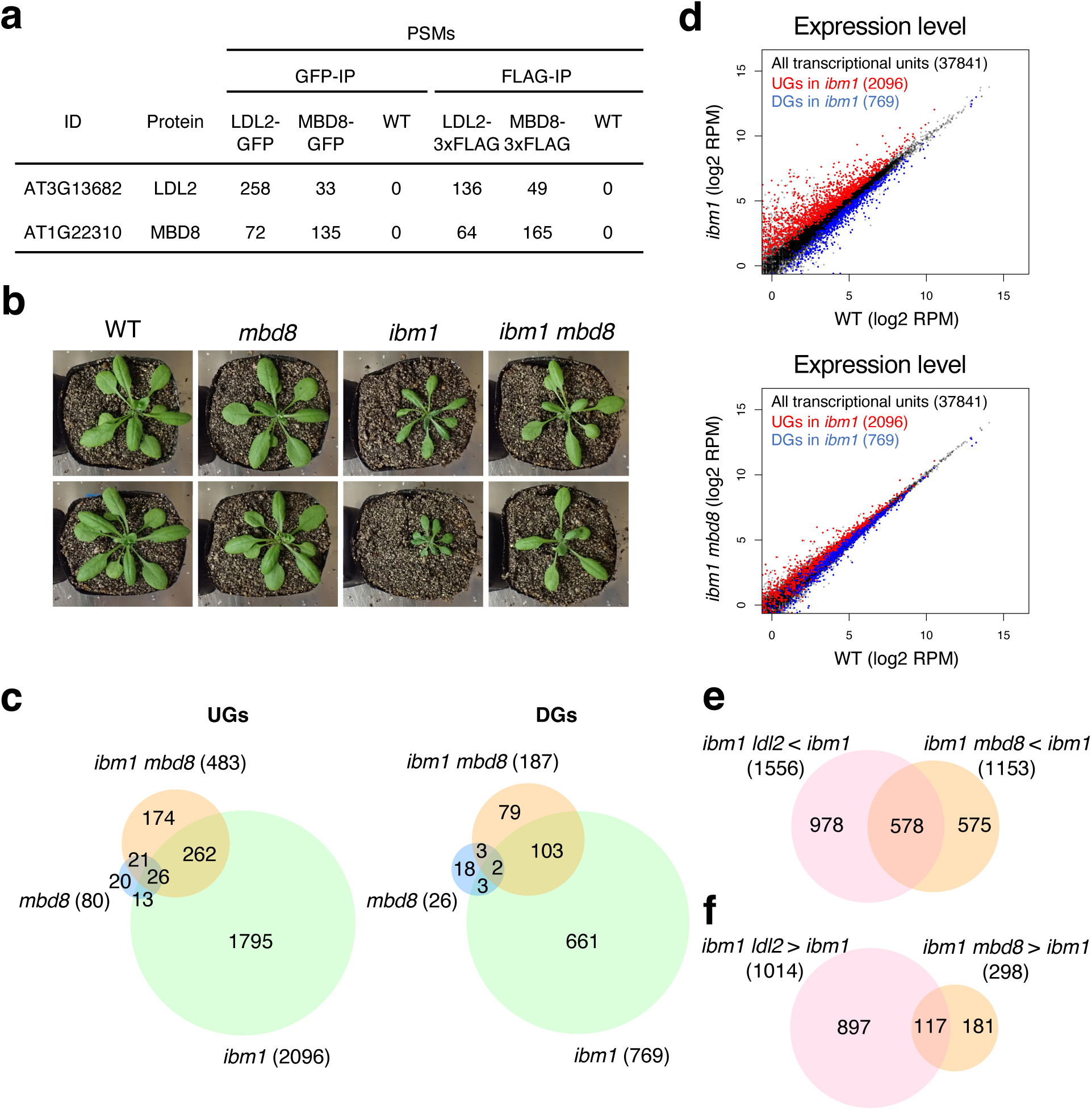
MBD8 binds to LDL2 and mediates the developmental defects of in the *ibm1* mutant. **a.** Epitope-tagged LDL2 and MBD8 proteins were immunoprecipitated and putative interacting proteins were identified by mass spectrometry. Peptide-spectrum match (PSM) are shown. WT replicates are used as a negative control. **b.** Phenotype of each mutant. Progeny were obtained from self-crossed *ibm1* +/- *mbd8-3* +/- plants, resulting in segregants including WT, *mbd8-3*, *ibm1* and *ibm1 mbd8-3* . The phenotypes of their offspring are shown. **c.** Venn diagram of the up-regulated genes (UGs) and downregulated genes (DGs) in each. **d.** Scatter plots showing the comparisons of expression level between WT and the *ibm1* (upper panel) or *ibm1 mbd8* (bottom panel) mutant. Red and blue dots representing UGs and DGs, respectively in *ibm1* shown in c. Their dots are superimposed over black dots representing all transcriptional units. Value represents the number of log_2_RPM. **e.** Ven diagram representation of the significant overlaps between down-regulated genes in *ibm1 ldl2* compared to *ibm1* ^35^ and down-regulated genes in *ibm1 mbd8* compared to *ibm1* (P <2.2 × 10^-308^, hypergeometric test). **f.** Ven diagram representation of the significant overlaps between up-regulated genes in *ibm1 ldl2* compared to *ibm1* ^35^ and up-regulated genes in *ibm1 mbd8* compared to *ibm1* (P = 4.3 × 10^-104^, hypergeometric test).

Thousands of genes are known to be misregulated in the *ibm1* mutant, and this abnormal expression is suppressed by the *ldl2* mutation ^35^. To investigate whether MBD8 contributes to the regulation of gene expression in *ibm1*, we performed mRNA-seq using *mbd8*, *ibm1*, *ibm1 mbd8,* and the wild-type control (WT, Col-0). First, we confirmed that MBD8 was properly knocked out in the *mbd8-3* T-DNA mutant (**Extended Data** Fig. 2a). Importantly, LDL2 expression levels were not reduced in *mbd8-3*, indicating that MBD8 likely affects LDL2 function or recruitment, not its transcription (**Extended Data** Fig. 2a). There were a few differentially expressed genes (DEGs) in *mbd8* compared with WT (80 upregulated genes (UGs) and 26 downregulated genes (DGs)) (**Fig. 1c**). Consistent with previous studies, thousands of genes were significantly misregulated in *ibm1* (2096 UGs and 769 DGs) ^35^. The number of misregulated genes was reduced in *ibm1 mbd8* (483 UGs and 187 DGs) (**Fig. 1c, d**). This result aligns with the WT-like developmental phenotypes of *ibm1 mbd8* (**Fig. 1b**). In both *mbd8* and *ibm1*, the majority of misregulated genes were protein-coding genes rather than TEs (**Extended Data** Fig. 2b). The genes with altered mRNA levels in *ibm1 mbd8* compared with *ibm1* significantly overlapped with the genes with altered mRNA levels in *ibm1 ldl2* (**Fig. 1e, f**). Taken together, these results suggest that MBD8 functions in the same pathway as LDL2 to regulate gene expression and development.

### MBD8 functions together with LDL2 to remove H3K4me1 from genes that accumulate H3K9me2

The *ldl2* mutation suppresses H3K4me1 reduction without affecting H3K9me2 or mCH levels in *ibm1* ^35^. To examine whether the *mbd8* mutation affects chromatin modifications as the *ldl2* mutation does, we performed whole-genome DNA methylation analyses (EM-seq) and ChIP-seq analyses for H3K4me1 and H3K9me2. In the *ibm1* mutant, mCH accumulated in protein-coding genes but not in TE genes (**Extended Data** Fig. 3a, b). This accumulation of mCH was also observed in *ibm1 mbd8* (**Extended Data** Fig. 3a, b). Consistent with the mCH levels, H3K9me2 similarly accumulated in *ibm1* and *ibm1 mbd8* (**Fig. 2a**), suggesting that the *mbd8* mutation affects downstream of H3K9me2/mCH rather than the silencing marks themselves.

**Figure 2.**
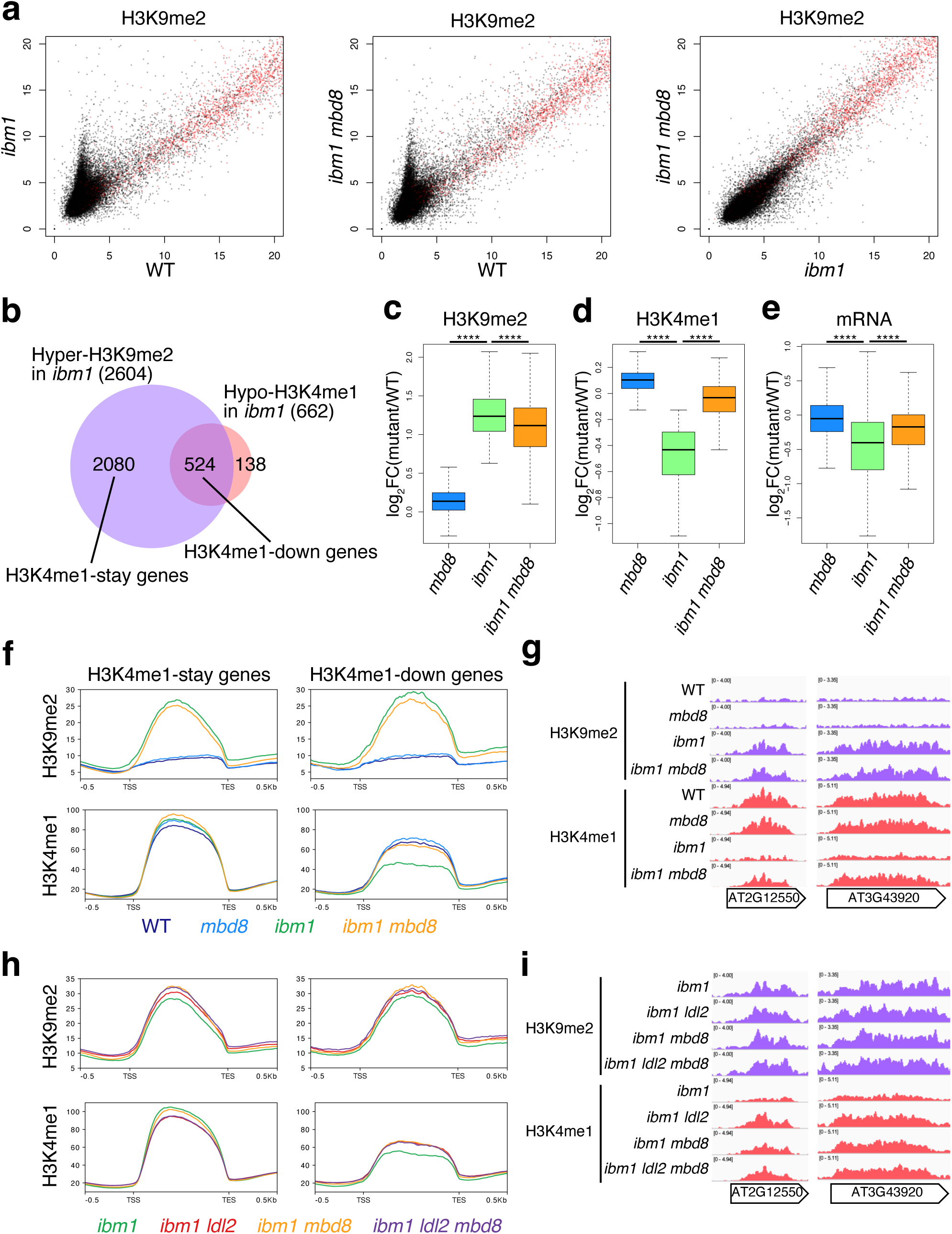
The *mbd8* mutation exhibited a similar suppression of H3K4me1 loss to that of the *ldl2* mutation. **a.** H3K9me2 levels in *ibm1* (left) or *ibm1 mbd8* (middle) compared to WT, and H3K9me2 levels in *ibm1 mbd8* compared to *ibm1* (right). Each dot represents the square root of RPKM within each transcription unit (black, protein-coding genes; red, TE genes). **b.** Venn diagram of the Hyper-H3K9me2 genes and Hypo-H3K4me1 genes in *ibm1*. **c, d.** Boxplots showing changes in H3K9me2 levels (**c**) and changes in H3K4me1 (**d**). Y-axis represents log_2_ fold change in RPKM values (mutant/WT). H3K4me1-down genes were analyzed, excluding genes shorter than 1.0 kb (total 524 genes). *P < 0.05; **P < 0.01; ***P < 0.001; ****P < 0.0001, Welch’s two-sample t-test. **e.** Boxplots showing changes in mRNA levels. Y-axis represents log_2_ fold change in RPM values (mutant/WT). H3K4me1-down genes were analyzed, excluding shorter than 1.0 kb or genes for which no transcription (total 470 genes). **f.** Averaged profiles of H3K9me2 and H3K4me1 around H3K4me1-stay genes and H3K4me1-down genes in WT, *mbd8, ibm1, ibm1 mbd8*. **g.** Genome browser views of H3K9me2 and H3K4me1 around H3K4me1-down genes in WT, *mbd8, ibm1, ibm1 mbd8*. **h.** Averaged profiles of H3K9me2 and H3K4me1 around H3K4me1-stay genes and H3K4me1-down genes in *ibm1, ibm1 ldl2, ibm1 mbd8, and ibm1 ldl2 mbd8.* **i.** Genome browser views of H3K9me2 and H3K4me1 around H3K4me1-down genes in *ibm1, ibm1 ldl2, ibm1 mbd8, and ibm1 ldl2 mbd8*.

To investigate the effect of the *mbd8* mutation on H3K4me1, we focused on hypo-H3K4me1 genes among the hyper-H3K9me2 genes whose H3K4me1 levels are reduced in the *ibm1* mutant. These genes (H3K4me1-down genes) are considered to be LDL2-target genes (**Fig. 2b, Extended Data** Fig. 4a, b). In *ibm1*, H3K4me1 levels were decreased in association with the increase in H3K9me2 (**Fig. 2c, d**). Although the *mbd8* single mutation did not affect H3K4me1 levels, H3K4me1 levels were restored in the *ibm1 mbd8* double mutant compared with those in the *ibm1* mutant (**Fig. 2d**); similarly, the mRNA levels were also restored in *ibm1 mbd8* (**Fig. 2e**). This suggests that the reduction of H3K4me1 in *ibm1* depends on MBD8 function. The *mbd8* mutations had effects on chromatin modifications similar to those of the *ldl2* mutation; MBD8 seems to function in H3K4me1 removal downstream of H3K9me2/mCH. The effect of *mbd8* on H3K4me1 was limited to H3K4me1-down genes, with no effect on the H3K4me1 levels of H3K4me1-stay genes, which did not exhibit a decrease in H3K4me1 in the *ibm1* mutant (**Fig. 2b, f, g**). To determine whether MBD8 and LDL2 function in the same pathway, we generated an *ibm1 ldl2 mbd8* triple mutant. The *ibm1 ldl2 mbd8* triple mutant showed comparable H3K4me1 recovery to that of the *ibm1 ldl2* and *ibm1 mbd8* double mutants (**Fig. 2h, i**), suggesting that MBD8 and LDL2 work through the same pathway for H3K4me1 removal. Considering that LDL2 is a putative histone demethylase for H3K4me1, MBD8 is likely to support H3K4me1 demethylation by LDL2.

### MBD8 binds both methylated and unmethylated DNA of AT-rich sequences in vitro

The next question we addressed is how MBD8 affects LDL2 function. First, we speculated that MBD8 binds to mCH because MBD8 contains a methyl-CpG-binding domain (MBD) and CHG/CHH methylation accumulates in the *ibm1* mutant. To test the binding ability of MBD8 to mC, we performed an electrophoretic mobility shift assay (EMSA) using the recombinant MBD8 protein. MBD8 proteins were incubated with 50 bp double-stranded DNA that contained three methylated cytosines. We designed DNA sequences containing each of the three contexts: CG, CHG, and CHH. Additionally, we prepared DNA with both strands methylated, one strand methylated (hemimethylated), and both strands unmethylated. MBD8 bound to all types of DNA regardless of sequence context or methylation (**Fig. 3a, Extended Data** Fig. 5). That result suggests that MBD8 does not show binding specificity toward mC. Since EMSA can assess binding affinity only for a certain DNA sequence, we performed DNA affinity purification sequencing (DAP-seq) to examine binding affinity across the entire genome sequence with natural mC. We incubated Halo-tag (HALO-STOP) or Halo-tagged MBD8 (HALO-MBD8) recombinant proteins with genomic DNA extracted from WT plants or *ibm1* mutant plants. We also used Halo-tagged MBD5 (HALO-MBD5) as a positive control, as MBD5 is known to exhibit a binding preference for mCG over unmethylated CG ^24^. As expected, MBD5 DAP-seq enrichment strongly correlated with mCG density (**Fig. 3b, Extended Data** Fig. 6a). In contrast, MBD8 did not show a binding preference for mC (**Fig. 3b, Extended Data** Fig. 6b), which is consistent with the EMSA results. Although mCH ectopically accumulated in the gene body in the *ibm1* mutant, there was no difference in the DAP-seq signals between DNA from the WT and the *ibm1* mutant (**Fig. 3b, c**). However, we observed a strong genome-wide correlation between MBD8 DAP-seq enrichment and AT density of the DNA (**Fig. 3d**). This result suggests that MBD8 does not preferentially bind to methylated DNA but binds to AT-rich DNA in vitro.

**Figure 3.**
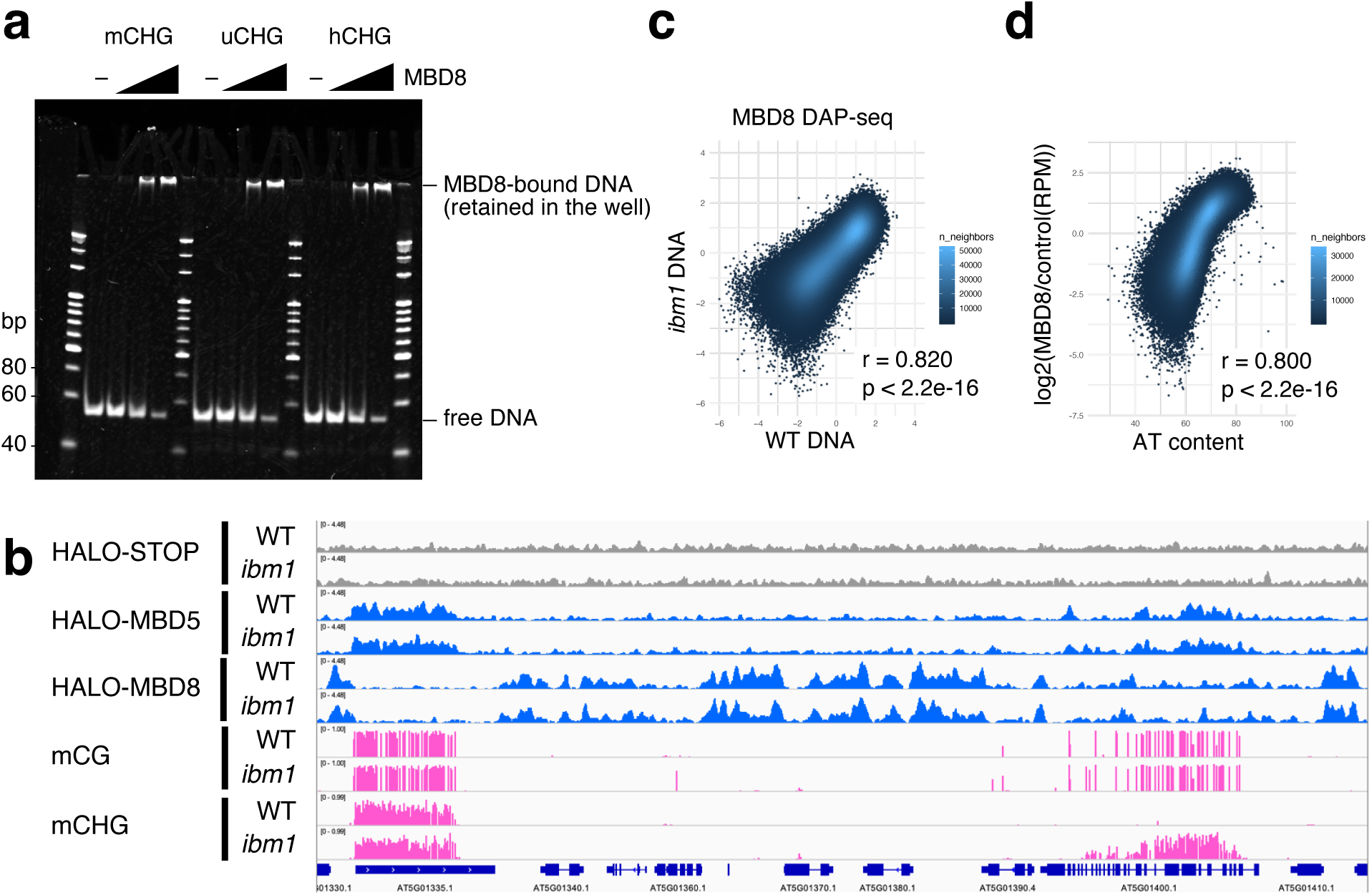
MBD8 did not show binding specificity towards mC and H3K9me2 in vitro. **a.** EMSA showing the binding of MBD8 to DNA that contains CHG sites. **b.** Genome browser views showing DAP-seq signals and DNA methylation levels in WT and *ibm1*. **c.** Scatterplots showing the correlation of MBD8 DAP-seq signals using DNA in WT and *ibm1*. Each dot represents a 400 bp bin. DAP-seq signals are normalized to the negative control (log_2_(HALO-MBD8/HALO-STOP)). **d.** Scatterplots showing the correlation of MBD8 DAP-seq signals and AT content in WT. Each dot represents a 400 bp bin. Bins for which no signal was detected or that had no AT were excluded.

In addition to mCH, H3K9me2 accumulates in the gene body of *ibm1*; thus, we next tested whether MBD8 binds to H3K9me2. We probed His-tagged recombinant proteins with a histone peptide array. As a control, we first used CMT2, which is known to bind H3K9me, and detected the preferential binding of CMT2 to peptides containing H3K9me (**Extended Data** Fig. 7a), which was consistent with the results of a previous study ^13^. However, we did not detect the binding of MBD8 to H3K9me but detected binding to H4K16ac and H4K20ac (**Extended Data** Fig. 7b). The affinity of MBD9 for acetylated H4 has been reported in previous studies ^28,29^, raising the possibility that paralogs share a similar affinity.

### MBD8 positively regulates LDL2 protein stability

Since the mRNA levels of LDL2 were not reduced in *mbd8* (**Extended Data** Fig. 2a), we next tested the possibility that MBD8 regulates LDL2 protein levels. FLAG-tagged LDL2 transgenic plants were crossed with *mbd8-3* mutant plants, and LDL2 protein levels were assessed by western blotting. Indeed, LDL2 protein levels were significantly reduced in the *mbd8* mutant background (**Fig. 4a, b**). We also found that the proteasome inhibitor MG132 restored the LDL2 protein level in *mbd8* to close to that in WT (**Fig. 4b, c**), suggesting that MBD8 positively regulates LDL2 protein stability by protecting LDL2 from proteasomal degradation. These results account for the dependence of LDL2 function on MBD8.

**Figure 4.**
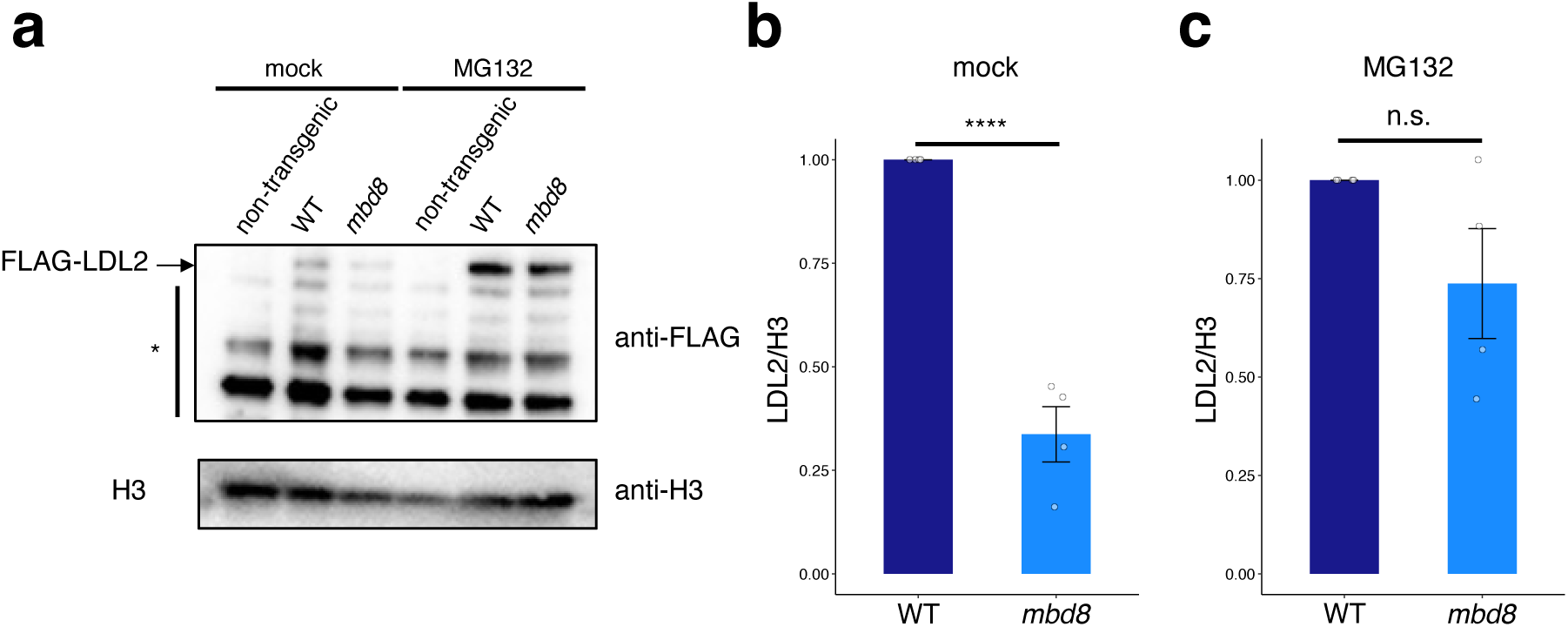
MBD8 positively regulates LDL2 protein stability. **a.** Western blot analysis showing the amount of 3xFLAG-LDL2 protein in WT and *mbd8* mutant backgrounds under mock and MG132 treatment conditions. The top panels of 3xFLAG-LDL2 were blotted with anti-FLAG, and the bottom panels were blotted with anti-H3 as loading controls. The asterisk indicates non-specific bands. **b, c.** Relative LDL2 protein levels under mock (**b**) and MG132 (**c**) treatment conditions as determined by Western blot. LDL2 protein signals were normalized to H3 and further standardized so that the protein level under mock conditions is set to 1. Means and SD for four technical replicates are shown. ****P < 0.0001, Welch’s two-sample t-test. n.s., not significant.

### The conserved N-terminal region in MBD8 is required for its interaction with LDL2

To determine which domain of MBD8 is required for the interaction with LDL2, we mapped the binding site of MBD8 on LDL2 using a series of truncated MBD8 proteins (**Fig. 5a**). We coexpressed recombinant Halo-tagged MBD8 and Strep-tagged LDL2 in *E. coli* and performed pull-down assays. While full-length MBD8 bound to LDL2, the deletion of the N-terminal 173 amino acid of MBD8 (ΔN-terminal) disrupted the interaction with LDL2 (**Fig. 5b**); conversely, the N-terminal 173 amino acid of MBD8 (ΔC-terminal) was sufficient for the interaction (**Fig. 5b**). Thus, the N-terminus of MBD8 is necessary and sufficient for the MBD8-LDL2 interaction.

**Figure 5.**
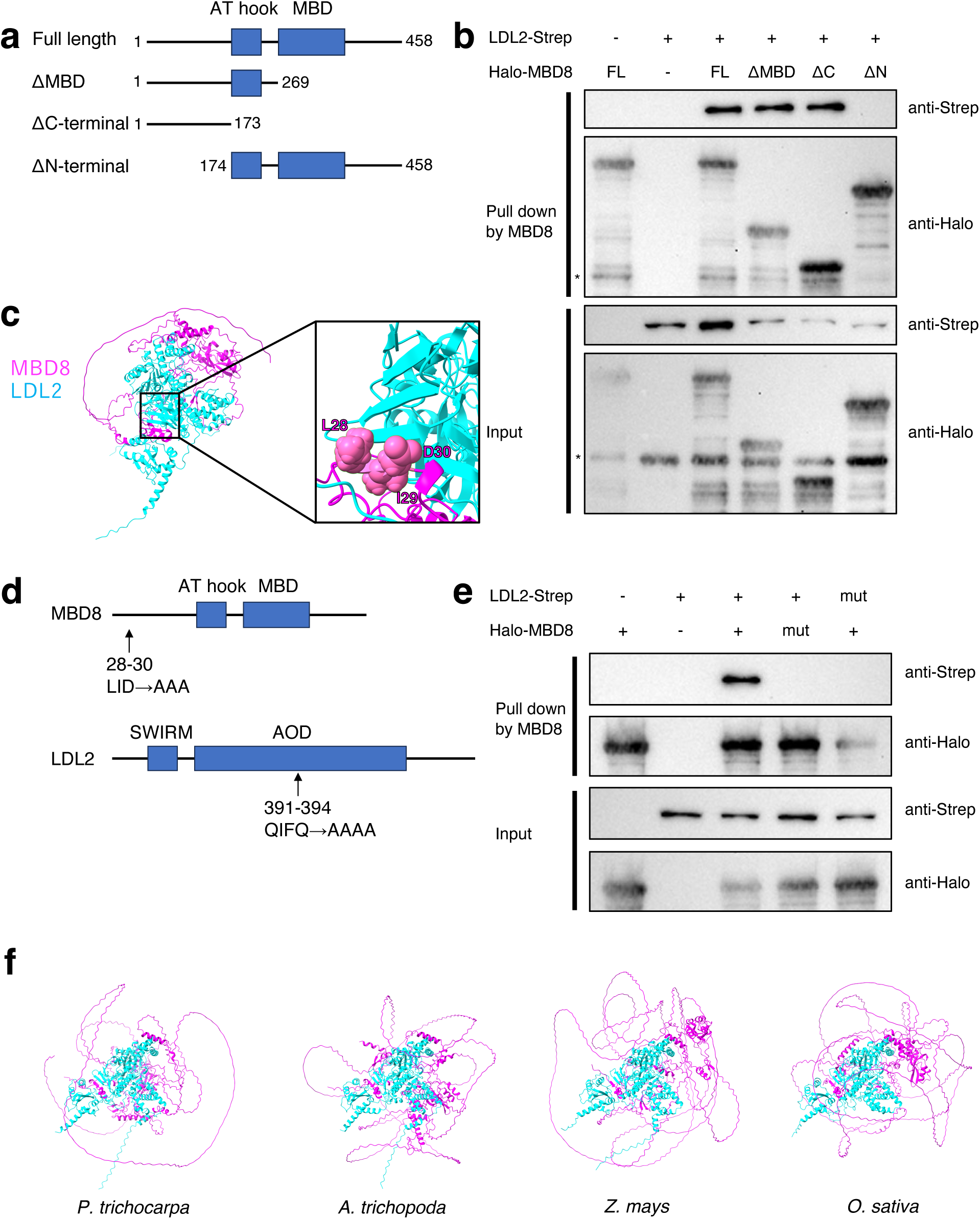
MBD8 N-terminal region is required for interaction with LDL2. **a.** Schematic representation of the truncated domain MBD8 protein. The full-length MBD8 protein (amino acids 1–458) contains two putative functional domains: AT hook and MBD. The numbers beside the diagram indicate the amino acid residues from the wild-type proteins. A Halo-tag at the N-terminus of all constructs is not shown. **b.** Pull-down assays testing the binding between LDL2 and each of the series of truncated MBD8 proteins. Bottom panels show input samples, and upper panels show proteins purified with MBD8. The asterisks indicate non-specific bands. **c.** MBD8-LDL2 complex structure predicted by Alphafold2. Structure zoom-in highlighting amino acid residues forming a beta-sheet structure between MBD8 and LDL2. **d.** Schematic representation of alanine substitution mutant constructs of MBD8 and LDL2 protein. **e.** Pull-down assays using mutated LDL2 protein and mutated MBD8 proteins. **f.** MBD8-LDL2 complex structure in four different plant species predicted by Alphafold3.

To further elucidate the binding modes of these proteins, we used AlphaFold-Multimer prediction. Interestingly, amino acid residues 28 to 30 (LID) of MBD8, which were disordered in the monomer structure prediction, formed a beta-sheet structure with amino acid residues 391 to 395 (QIFQA) of LDL2 in the multimer structure prediction (**Fig. 5c, Extended Data** Fig. 8a-c). This suggests that those amino acids play a role in the MBD8-LDL2 interaction. To test this possibility, we substituted those amino acids in MBD8 and LDL2 with alanine (**Fig. 5d**). Pull-down assays revealed that both substitutions in MBD8 and LDL2 disrupted the interaction (**Fig. 5e**), suggesting that the LID motif of MBD8 and the QIFQA motif of LDL2 are necessary for the MBD8-LDL2 interaction. The long N-terminal region of MBD8 is unique to AtMBD family proteins ^19^, and the QIFQA motif of LDL2 is also unique to AtLSD1 family proteins (**Extended Data** Fig. 9). Thus, it is expected that they bind to each other specifically. We found that these residues of MBD8 are conserved in land plants (**Extended Data** Fig. 10), and the MBD8 and LDL2 pairs in other plants are also predicted to form complexes, as determined by AlphaFold3 (**Fig. 5f**) ^38^, suggesting that the MBD8LJLDL2 interaction may also be conserved in land plants.

## Discussion

LDL2 is known to remove H3K4me1 and function in gene silencing downstream of H3K9me2/mCH ^35^. However, how LDL2 functions in H3K9me2/mCH-marked genes is unclear. In this study, we screened for LDL2-binding proteins and identified a methyl CpG-binding domain family protein, MBD8. Genetic analysis revealed that MBD8 and LDL2 function in the same pathway. The *mbd8* mutation did not affect H3K9me2/mC levels but affected H3K4me1 levels in H3K9me2/mCH-marked genes, suggesting that MBD8 functions downstream of H3K9me2/mC and is required for H3K4me1 demethylation by LDL2. Although we expected that MBD8 would bind to H3K9me2 or mCH, it exhibited a binding affinity for AT-rich DNA sequences and H4 acetylation. As these features are not specific to LDL2 target genes, it remains unclear whether MBD8 determines the chromatin localization of LDL2. As MBD8 does not retain the conserved amino acid residues required for binding to methylated DNA ^18^, it is not surprising that MBD8 did not show binding specificity toward mC, as determined by EMSAs and DAP-seq. However, MBD1 and MBD2, which are known to be localized in chromatin with high levels of mCG in vivo, do not show any signals in DAP-seq ^26^, suggesting that additional factors are necessary for their specific chromatin localization in vivo. Thus, it is reasonable to suspect that MBD8 may bind mC in vivo. To address these possibilities, we employed ChIP-seq for MBD8 using tagged MBD8 transgenic plants, but we did not observe any significant peaks where mC exists in WT and *ibm1* plants. LDL2 also did not show any ChIP-seq signal, suggesting that MBD8/LDL2 binding to chromatin may be weak or transient. Elucidating the chromatin localization of MBD8/LDL2 by new techniques will provide further insight into LDL2 regulation by MBD8. Interestingly, we found that MBD8 positively regulates LDL2 protein stability. That finding suggests a possible dynamic regulation by which transcription is repressed through the modulation of LDL2 protein levels. In animals, the protein stability of LSD1, an LDL2 homolog, is regulated by the cofactor CoREST ^39^. Considering that plants do not have CoREST homologs ^40^, MBD8 possibly functions as an alternative to CoREST.

The biological role of the MBD8/LDL2 pathway is still enigmatic. This pathway has a significant phenotypic impact but is normally masked by the H3K9 demethylase IBM1 ^32–35^. Examination of other components of the pathways to write and read H3K9me may clarify the view of this enigmatic pathway conserved among plants.

## Materials and methods

### Plant materials and growth conditions

*A. thaliana* strain Columbia-0 (Col-0) was used as the wild type (WT). The *ibm1-4* (SALK_035608) ^32^ and *ldl2-1* (SALK_135831) ^41^ strains have been previously described. We used CS881802 as *mbd8-3*. CRISPR-Cas9 mutants for MBD8 were generated with the pPLV01-pAT2S3::GFP-pRPS5::Cas9 vector, which contains two guide RNAs for the MBD8 gene. The vector was transformed into *ibm1* heterozygote mutant plants, and T1 plants were selected by GFP fluorescence. T2 plants in which the transgene had segregated out were selected. GFP-tagged and FLAG-tagged MBD8 lines were generated by cloning the genes from genomic DNA into the pGreenII 0179 vector. The MBD8 DNA fragment from its endogenous promoter to the last base before the stop codon was cloned and fused in-frame with a C-terminal tag. pLDL2::LDL2-FLAG plants have been previously described ^35^. Plants were grown on Murashige and Skoog (MS) medium supplemented with 1% sucrose and solidified with Bacto agar for 14 days under long-day (16 h light/8 h darkness) photoperiods at 22°C. For LDL2 protein stability assays, plants were cultivated for 14 days on MS plates and then transferred to liquid MS medium supplemented with 50 µM MG132 (1000x diluted from 50 mM MG132 stock solution in DMSO) or DMSO as a control. Twenty-four hours after transfer, the plants were collected and frozen with liquid nitrogen for nuclear isolation.

### Immunoprecipitation mass spectrometry (IP-MS)

The IPLJMS experiments were performed as previously described ^42^ with minor variations. We used approximately 2 g of seedlings of FLAG- or GFP-tagged lines and of nontransgenic plants as a negative control. The samples were ground in liquid nitrogen and resuspended in 10 mL extraction buffer [25 mM HEPES-KOH pH 7.4, 100 mM NaCl, 2 mM MgCl2, 10% glycerol, 0.05% IGEPAL CA-630, 1 mM DTT, 5 mM EGTA, 1 tablet/50 mL cOmplete]. Then, 30 µL of 1 M MgCl2 (final concentration of 5 mM) and 2 µL of 250 U/µL benzonase (final concentration of 50 U/mL) were added, and the mixture was incubated for 30 min at 4°C on a rotator. After centrifugation at 4,000 × *g* at 4°C, the suspension was filtered through a 40 µm nylon cell strainer. Dynabeads protein G beads (40 μL) were washed with PBST buffer (phosphate-buffered saline (PBS) with 0.1% Tween 20) twice and resuspended in 200 μL of PBST buffer. Anti-FLAG (F1804; SIGMA) or anti-GFP (ab290; Abcam) antibody (10 μg) was added to the beads, followed by incubation for at least 2 h on a rotator at 4°C to form antibody-bead complexes. The antibody-bead complexes were washed with PBST twice and resuspended in the extraction buffer. The resuspended antibody-bead complex was incubated with the protein extract for 2 h at 4°C on a rotator. The sample was centrifuged (2000 × *g*, 15 min) to pellet the beads, and the supernatant was removed. The beads were resuspended in 1 mL of extraction buffer, transferred to a 1.5 mL tube, and pelleted with a magnetic rack, after which the supernatant was removed. The wash step was repeated two additional times, followed by protein digestion.

Protein and nano LCLJMS/MS data were acquired by the Matsubayashi laboratory at Nagoya University, Nagoya, following the same protocol as described previously^43^.

### mRNA-seq

Total RNA was isolated from the aerial part of one seedling using an RNeasy Plant Mini Kit (Qiagen). Libraries for mRNA-seq were constructed using a KAPA Stranded RNA-seq Library Preparation Kit, with 800 ng of RNA as input, following the manufacturer’s instructions. Three independent biological replicates were analyzed for each genotype.

Sequencing reads were evaluated for quality and trimmed for adapter sequences and low-quality bases using Fastp (v0.20.1) ^44^. Trimmed reads were mapped to the Arabidopsis TAIR10 reference genome using STAR (v2.7.9a) ^45^ with --outFilterType BySJout --alignSJoverhangMin 8 --alignSJDBoverhangMin 1 --quantMode GeneCounts parameters.

### Enzymatic methyl-seq

Whole-genome DNA methylation sequencing was performed as previously described^46^. Genomic DNA was extracted from seedlings using Nucleon PhytoPure (Cytiva). Approximately 500 ng of DNA was fragmented into 400–600 bp fragments using a Covaris S220 focused ultrasonicator (Covaris) and size-selected with SPRIselect beads (Beckman Coulter). EM-seq libraries were prepared from 30 ng of fragmented DNA using an NEBNext Enzymatic Methyl-seq Kit (NEB) following the manufacturer’s instructions. Two independent biological replicates were analyzed per genotype. Libraries were 150-bp paired-end sequenced using HiSeq X (Illumina). The sequencing reads were trimmed for adapter sequences and low-quality bases using Trimmomatic ^47^. The trimmed reads were aligned to the Arabidopsis TAIR10 reference genome using Bismark (v0.10.1) ^48^ with -n 1 - l 20 - e 90 parameters. Deduplication and methylation extraction were also performed using Bismark. Methylation levels were calculated as the ratio of methylated to total cytosines within each transcription unit using the “map” function in BEDTools ^49,50^. Metaplots were generated using deepTools ^51^.

### ChIP-seq

Chromatin immunoprecipitation sequencing (ChIP-seq) was carried out as previously described ^52^ with modifications. A total of 0.5∼0.6 g of aerial parts from seedlings was used for each ChIP sample. The samples were ground in liquid nitrogen and crosslinked for 10 min at room temperature in nuclei isolation buffer [10 mM HEPES pH 7.5, 1 M sucrose, 5 mM KCl, 5 mM MgCl2, 5 mM EDTA, 0.1% b-mercaptoethanol, 0.6% Triton X-100, 1 tablet/50 ml cOmplete proteinase inhibitor and 1 mM Pefabloc SC (Roche)] supplemented with 1% formaldehyde. The crosslinking reaction was stopped with 130 mM glycine. The suspension was filtered through a 40 µm nylon cell strainer and pelleted by centrifugation at 3,000 × *g* at 4°C for 10 min. The pellet was resuspended in 300 µL of nuclei isolation buffer and layered on top of 500 µL of nuclei separation buffer [10 mM HEPES pH 7.6, 1 M sucrose, 5 mM KCl, 5 mM MgCl2, 5 mM EDTA pH 8.0, 15% Percoll] and pelleted by centrifugation at 3,000 × *g* for 10 min at 4°C. The nuclear pellet was resuspended in 900 µL of RIPA without Triton buffer [50 mM Tris–HCl pH 7.8, 150 mM NaCl, 1 mM EDTA, 0.1% SDS, 0.1% sodium deoxycholate, and cOmplete proteinase inhibitor]. Sonication was conducted using a Picoruptor (Diagenode) (6–8 cycles of 30 s on and 30 s off).

The sonicated chromatin was then centrifuged at 13,000 × *g* for 3 min, and 10 µL of 10% Triton X-100 was added to the supernatant, which was subsequently aliquoted. The chromatin mixture was incubated with 1–2 µg of antibodies overnight at 4°C. The antibodies used were anti-H3K4me1 (49--1003, Invitrogen), anti-H3K9me2 (MABI0317, MBL), and anti-H3 (ab1791; Abcam).

Then, the antibody-chromatin mixture was incubated for 2 h at 4°C with magnetic beads: Dynabeads M280 Sheep anti-mouse IgG in the case of MBL antibodies and Dynabeads Protein G in the case of the other antibodies. The beads were washed once with low-salt ChIP buffer [50 mM HEPES-KOH (pH 7.5), 150 mM NaCl, 1 mM EDTA, 1% Triton X-100, 0.1% sodium deoxycholate, and 0.1% SDS] containing cOmplete proteinase inhibitor cocktail, two times with high-salt ChIP buffer [50 mM HEPES-KOH (pH 7.5), 350 mM NaCl, 1 mM EDTA, 1% Triton X-100, 0.1% sodium deoxycholate, and 0.1% SDS], once with LNDET buffer [0.25 M LiCl, 1% IGEPAL, 1% sodium deoxycholate, 1 mM EDTA, and 10 mM Tris pH 7.8], and once with TE buffer. The chromatin was eluted by adding ChIP elution buffer [50 mM Tris-HCl (pH 7.5), 10 mM EDTA, and 1% SDS] and incubating at 65°C for 15 min. Subsequently, 5 μl of proteinase K (20 mg/ml; Thermo Fisher Scientific) was added to the sample, followed by incubation at 55°C overnight. The immunoprecipitated DNA was purified using a Monarch PCR & DNA Cleanup Kit (New England Biolabs). The libraries for Illumina sequencing were constructed using a ThruPLEX DNA-Seq Kit (Clontech) and purified using SPRIselect Beads (Beckman Coulter). Sequencing was performed using a HiSeq X or NovaSeq X Plus sequencer (Illumina).

Sequencing reads were trimmed using Fastp (v0.20.1) ^44^ and then mapped to the Arabidopsis TAIR10 reference genome using Bowtie2 (v2.4.4) ^53^. The read count for each transcription unit was calculated using the coverage function in BEDTools ^49^, and then, the reads per kilobase per million mapped reads (RPKM) values were calculated. Metaplots were generated using deepTools ^51^. The mapping results were visualized with IGV ^54^ using TDF files generated by the igvtools (v2.5.3) count function ^55^.

### Protein purification

MBD8 expression and purification were performed using a previously described method ^56,57^. Briefly, MBD8 proteins (full-length) were expressed in BL21 (DE3) *E. coli.* containing a minor tRNA expression vector (Codon (+) RIL) from a pET-15b plasmid containing an N-terminal 6xHis tag and a Halo tag. After purification using Ni-NTA resin, MBD8 was subjected to a RESOURCE Q anion exchange column (Cytiva), collected from the flow-through fractions, and then purified on a RESOURCE S cation exchange column (Cytiva). MBD8 protein was further purified on a Superdex 200 gel filtration column using storage buffer (20 mM Tris-HCl (pH 7.5), 150 mM NaCl, 10% glycerol, and 2 mM 2-mercaptoethanol). The process for CMT2 was the same as that for MBD8, except that N-terminally truncated CMT2 (residues 504–1,295) was cloned ^13^.

### MBD8-DNA binding assay

Oligonucleotides (**Supplementary Table 1**) were obtained and annealed in annealing buffer [10 mM Tris-HCl (pH 7.5), 100 mM NaCl, 1 mM EDTA] by heating and cooling. The 50 bp DNA (0.2 μM) was mixed with MBD8 (0, 0.2, 0.8, and 1.6 μM) in a total volume of 10 μl of reaction buffer [20 mM Tris-HCl (pH 7.5), 150 mM NaCl, 10% glycerol, and 2 mM 2-mercaptoethanol]. The samples were incubated at 25°C for 30 min and then analyzed by 8% nondenaturing PAGE in 0.5×TBE. The gel was stained with SYBR Green I solution, and the DNA was visualized with an iBright Imaging System (Thermo Fisher Scientific).

### DAP-seq

DNA affinity purification sequencing (DAP-seq) was performed as described previously ^58^ with slight modifications. First, the MBD5 and MBD8 genes were cloned from cDNA into the pENTR/D-TOPO vector (Thermo Fisher) from the ATG start codon until the stop codon. The genes were subsequently transferred into the pIX-HALO vector ^59^ using a Gateway LR Clonase II reaction. The pIX-STOP vector was used as a negative control. For DNA library preparation, genomic DNA was purified with Nucleon PhytoPure (Cytiva) from either WT (Col-0) seedlings or *ibm1-4* seedlings. The DNA was diluted with water into 125 µl aliquots containing 5 µg of DNA each and sheared with a Covaris S220 focused ultrasonicator (Covaris) to an average fragment size of 200 bp (peak incident power = 175, duty factor = 10%, cycles per burst = 200, and treatment time = 140 s). After ethanol precipitation, 1 µg of sheared DNA was subjected to library construction using an NEBNext UltraII DNA Library Prep Kit for Illumina (E7645, NEB). For recombinant protein expression, 2000 ng of plasmid DNA and the TNT SP6 High-Yield Wheat Germ Protein Expression System (Promega) were used. The protein was incubated with 50 ng of genomic DNA. The pulled-down DNA was amplified, purified, and sequenced.

Subsequent analysis was the same as ChIP-seq, except that the genome was divided into 400 bp bins, and the read count for each bin was calculated using the coverage function in BEDTools ^49^

### Histone peptide array assay

A total of 30 μg of recombinant CMT2 or MBD8 protein was screened on a MODified Histone Peptide Array slide following the manufacturer’s instructions (Active Motif). Briefly, the array was initially immersed in blocking solution (TBST + 5% milk) at room temperature for 30 min. After three washes with TBST, the array was incubated with the target protein in 5 mL of binding buffer (50 mM Tris-HCl pH 7.5, 150 mM NaCl, 5% glycerol, 2 mM DTT, 0.4% BSA) for 2 h at room temperature. Following another three TBST washes, the array was incubated overnight at 4°C with an anti-His-tag primary antibody (M089-3, MBL) diluted in blocking solution. After three TBST washes, the array was subsequently incubated with a secondary antibody (NA931, Cytiva) diluted in blocking solution for 1 h at room temperature. After a final set of three TBST washes, the signal was detected using Clarity Max ECL Western Blotting Substrates (Bio-Rad) and an iBright Imager (Thermo Fisher Scientific).

### Pull-down assay

As described for protein purification, MBD8 proteins were cloned and inserted into a pET15b plasmid containing an N-terminal 6xHis tag and a Halo tag. LDL2 proteins were cloned and inserted into a pET-28a plasmid, where the His-tag sequence was substituted with the Strep-tag II sequence at the C-terminus. Two vectors were introduced into BL21 (DE3) *E. coli* containing a minor tRNA expression vector (Codon (+) RIL), and colonies that had both vectors were selected on LB medium supplemented with ampicillin and kanamycin. The colonies were incubated in 200 mL of LB and grown at 25°C to an optical density of 0.6. Protein expression was induced with 0.5 mM IPTG, and the bacteria were subsequently grown overnight (>12 h) at 18°C. The bacteria were collected and lysed via sonication in buffer 1 [50 mM Tris-HCl pH 7.5, 500 mM NaCl, 10% glycerol, 1 mM phenylmethylsulfonyl fluoride, and 2 mM 2-mercaptoethanol]. The cell debris was removed by centrifugation, and the lysate (used as input) was mixed with Ni-NTA agarose resin (Qiagen) at 4°C for 1 h. After binding, the resin was washed with buffer 2 [50 mM Tris-HCl pH 7.5, 500 mM NaCl, 10% glycerol, 2 mM 2-mercaptoethanol, and 5 mM imidazole]. His-Halo-tagged MBD8 protein complexes were eluted by the addition of buffer 3 [50 mM Tris-HCl pH 7.5, 500 mM NaCl, 10% glycerol, 2 mM 2-mercaptoethanol, and 300 mM imidazole].

### Western blot

For the LDL2 protein stability assay, nuclei from 0.5 g of seedlings were isolated as described for ChIP-seq. The nuclear pellet was resuspended in 200 µL of SDS buffer and boiled for 10 min at 95°C. Proteins were separated by SDSLJPAGE using 4–20% Mini-PROTEAN^®^ TGX™ Precast Protein Gels (Bio-Rad) and then transferred to a PVDF membrane using a Trans-Blot Turbo and Trans-Blot transfer pack (Bio-Rad). The membrane was immersed in blocking solution (TBST + 5% milk) at room temperature for 30 min. After the blocking solution was removed, western blot analysis was performed using antibodies against FLAG (F1804, Sigma) as the primary antibody, anti-mouse IgG (NA931, Cytiva) as the secondary antibody, and Can Get Signal solution (TOYOBO). The signals were detected using Clarity Max ECL Western Blotting Substrates (Bio-Rad) and an iBright Imager (Thermo Fisher Scientific). For the pull-down assay, the pulled-down proteins were separated by 12% SDSLJPAGE and then transferred to a PVDF membrane using a Trans-Blot Turbo and Trans-Blot transfer pack (Bio-Rad). Western blot analysis was performed using an iBind Flex Western System (Thermo Fisher Scientific). Primary antibody (Strep (M211-3, MBL) and Halo (G9281, Promega)) and secondary antibody (anti-rabbit IgG HRP (NA934, Cytiva) and anti-mouse IgG HRP (NA931, Cytiva)) reactions were performed at room temperature for more than 2.5 h following the manufacturer’s protocol.

## Data availability

The data have been deposited with links to the BioProject accession number PRJNA1300776 in the NIH BioProject database.

## Acknowledgments

We thank all the Kakutani laboratory members. The pPLV01-pAT2S3::GFP-pRPS5::Cas9 vector was kindly provided by Dr. Taku Sasaki. The pIX-Halo-ccdB-CmR and pIX-Halo-STOP vectors were kindly provided by Prof. Taiji Kawakatsu and Prof. Joseph Ecker. Nano LCLJMS/MS and mapping were conducted by Prof. Yoshikatsu Matsubayashi and Dr. Saki Noda. The computations were partially performed on the NIG supercomputer at NIG, Japan. We also thank NASC/ABRC for providing the seeds. This work was supported by grants from the Human Frontier Science Program (HFSP) (RGP0025/2021) to T.K., Japan Science and Technology Agency (JST), CREST (nos. JPMJCR15O1 to T.K., and JPMJCR20S6 to A.O.), and the Japan Society for the Promotion of Science (JSPS) (nos. 21H04977, 23H00365 and 24K21268 to T.K., 20H05913, 23K23565, 24K21926, and 25K02252 to S.I., and 24H01351, 24K02077, 24K02002, 25K02258, 25H02544, and 25H01300 to A.O.).

## Author contributions

S.M., A.O., S.I., and T.K. conceived the study. S.M., J., Y.T., and M.H. performed the experiments. S.M. conducted the analysis and made the figures. S.M. wrote the manuscript, and S.M., A.O., S.I., and T.K. edited the manuscript.

---

**Extended Data Fig. 1. *mbd8* mutants’ alleles and *mbd8* mutants’ phenotype.**

**a.** Graphical representation of the *MBD8* gene showing the position of the mutations. The black boxes represent the exons. **b.** Phenotype of *ibm1* and *ibm1 mbd8-4* mutants.

**Extended Data Fig. 2. mRNA-seq analysis in each mutant.**

**a.** mRNA level (reads per million mapped reads; RPM) of *LDL2*, *IBM1* and *MBD8* genes in WT, *mbd8*, *ibm1*, and *ibm1 mbd8*. Means and SD for three biological replicates are shown. **b.** The numbers of up-regulated and down-regulated genes in each mutant.

**Extended Data Fig. 3. Whole-genome DNA methylation analysis in each mutant.**

**a,b.** Boxplots showing DNA methylation levels in each context (CG, CHG, and CHH; H can be A, T, or C) in all protein-coding genes (**a**), and all TE genes (**b**)

**Extended Data Fig. 4. Hyper-H3K9me2 genes and Hypo-H3K4me1 genes in *ibm1.***

**a.** The overlap of genes with increased H3K9me2 in *ibm1* compared to WT (H3K9me2 RPKM change (*ibm*1–WT) > 2) between two biological replicates. “Hyper-H3K9me2 genes in *ibm1*” are defined as the overlapping genes (n=2,604). **b.** The overlap of genes with decreased H3K4me1 in *ibm1* compared to WT (H3K4me1 RPKM change (*ibm*1–WT) < -2) between two biological replicates. “Hypo-H3K4me1 genes in *ibm1*” are defined as the overlapping genes (n=662).

**Extended Data Fig. 5. EMSA for DNA binding activity of MBD8 to CG and CHH sites.**

**a.** EMSA showing the binding of MBD8 to DNA that contains CG and CHH sites. We used oligonucleotides that were fully methylated (m), unmethylated (u), and hemimethylated (h).

**Extended Data Fig. 6. MBD5 shows binding specificity for mC, but MBD8 does not.**

**a,b.** Scatterplots showing the correlation of MBD5 DAP-seq signals (**a**) or MBD8 DAP-seq signals (**b**) and DNA methylation density (mC are normalised to mapped reads) in the indicated context. Each dot represents a 400 bp bin.

**Extended Data Fig. 7. Histone peptide array.**

**a.** CMT2 binding assay using a histone peptide array. Red and blue circles indicate peptides containing di-, and trimethylated H3K9 peptides, respectively. **b.** MBD8 binding assay. Peptides containing both H4K16ac and H4K20ac (green circles) are reproducibly bound to MBD8.

**Extended Data Fig. 8. MBD8 and LDL2 structure predicted by Alphafold2.**

**a.** MBD8-LDL2 complex structure predicted by Alphafold2. This is the same as (**Fig. 4c**). **b.** LDL2 monomer structure. **c.** MBD8 monomer structure.

**Extended Data Fig. 9. Amino acid alignment of LDL2 and its paralogs.**

**a.** In the alignment, positions conserved or highly similar among the proteins are colored in green for hydrophobic amino acids (VILM), blue for basic (KRH) amino acids, and pink for acidic (DE) amino acids. The asterisk symbol (*) refers to amino acid residues required for binding to MBD8. Protein names and domain limits (in parentheses) are indicated at the start of the sequences.

**Extended Data Fig. 10. Amino acid alignment of the N-terminal region of MBD8 and its orthologs.**

**a.** In the alignment, positions conserved or highly similar among the MBD8 proteins are colored in green for hydrophobic amino acids (VILM), blue for basic (KRH) amino acids, and pink for acidic (DE) amino acids. The asterisk symbol (*) refers to amino acid residues required for binding to LDL2. Host species and domain limits (in parentheses) are indicated at the start of the sequences. NCBI accession numbers are as follows: *Arabidopsis thaliana*: NP_173650.3, *Popilus trichocarpa*: KAI5597816.1, *Amborella trichopoda*: XP_020528307.1, *Zea mays*: NP_001123600.1, *Oryza sativa japonica*: XP_015611640.1.

**Supplementary Table 1.**
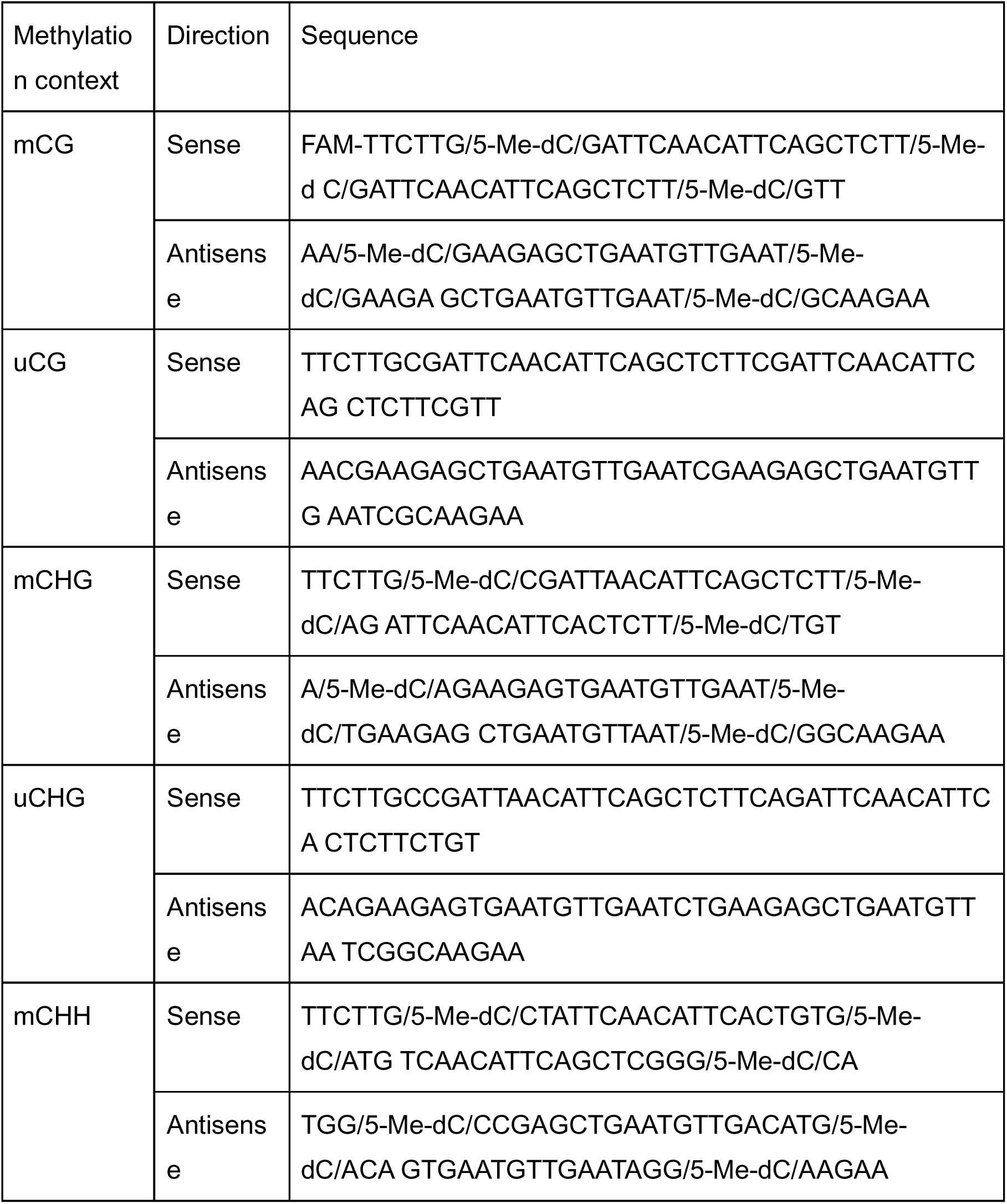
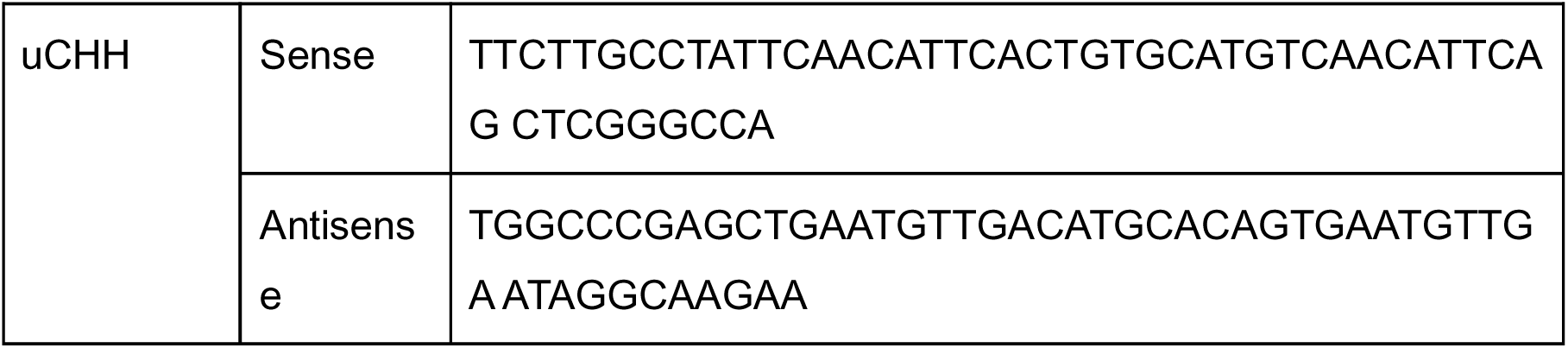
Oligonucleotides for MBD8-DNA binding assay.

