## Extended Data for "MBD8 is required for LDL2-mediated transcriptional repression downstream of H3K9me2 in *Arabidopsis*"

### Extended Data Fig.1

**a**

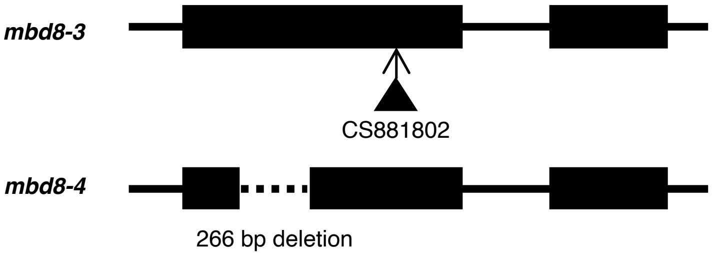


**b**

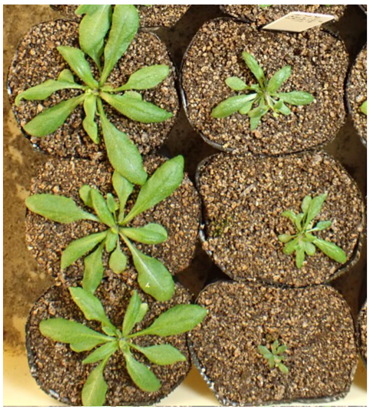


### Extended Data Fig.2

**a**

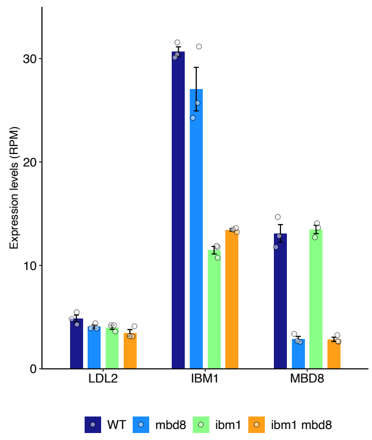


**b**

**UGs**

| *mbd8* | | *ibm1* | | *ibm1 mbd8* | |
|---|---|---|---|---|---|
| protein coding gene | 57 | protein coding gene | 1916 | protein coding gene | 387 |
| pseudo gene | 1 | pseudo gene | 18 | pseudo gene | 10 |
| TE | 7 | TE | 50 | TE | 42 |
| non-coding RNA | 11 | non-coding RNA | 101 | non-coding RNA | 33 |
| novel transcribed region | 4 | novel transcribed region | 11 | novel transcribed region | 11 |
| | 80 | | 2096 | | 483 |

**DGs**

| *mbd8* | | *ibm1* | | *ibm1 mbd8* | |
|---|---|---|---|---|---|
| protein coding gene | 22 | protein coding gene | 701 | protein coding gene | 163 |
| pseudo gene | 0 | pseudo gene | 11 | pseudo gene | 4 |
| TE | 1 | TE | 17 | TE | 6 |
| non-coding RNA | 2 | non-coding RNA | 38 | non-coding RNA | 11 |
| novel transcribed region | 1 | novel transcribed region | 2 | novel transcribed region | 3 |
| | 26 | | 769 | | 187 |

**Extended Data Fig.3**

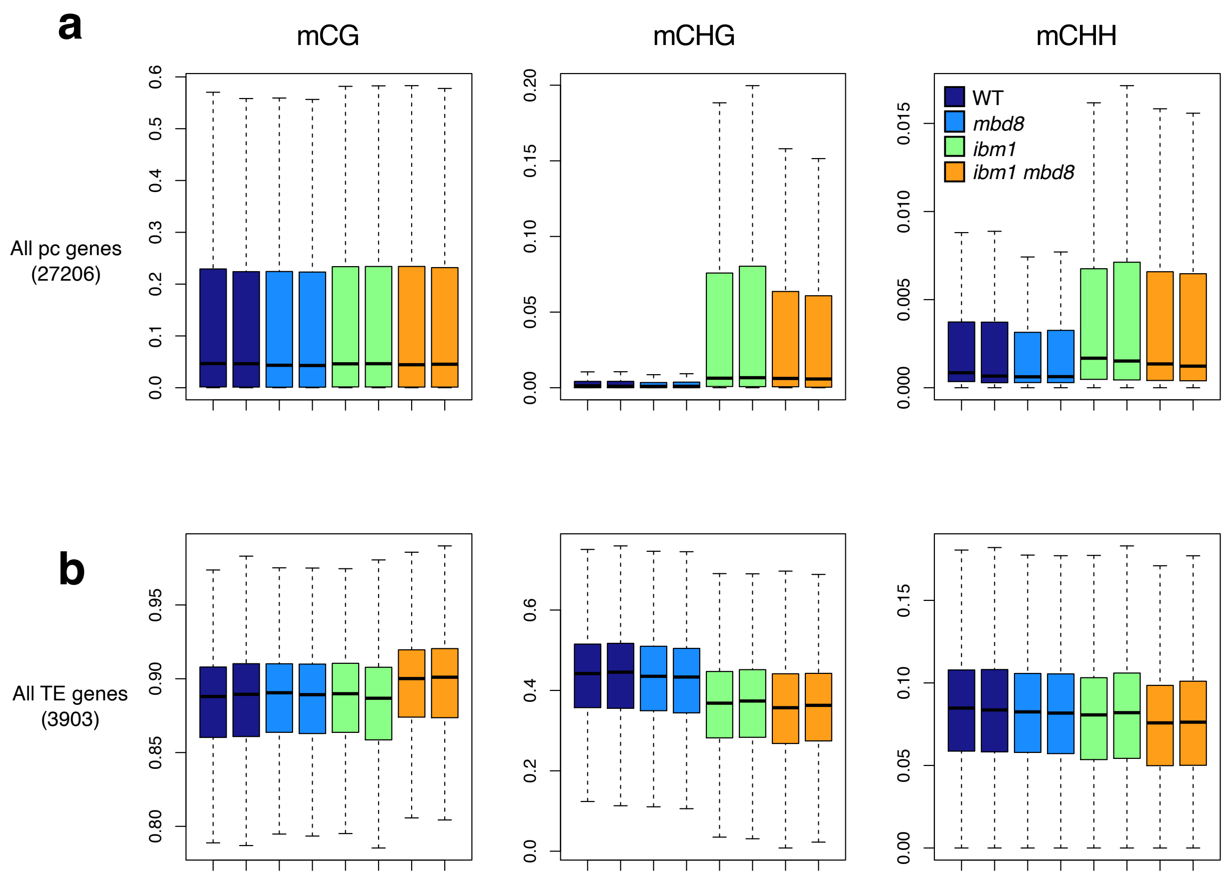


**Extended Data Fig.4**

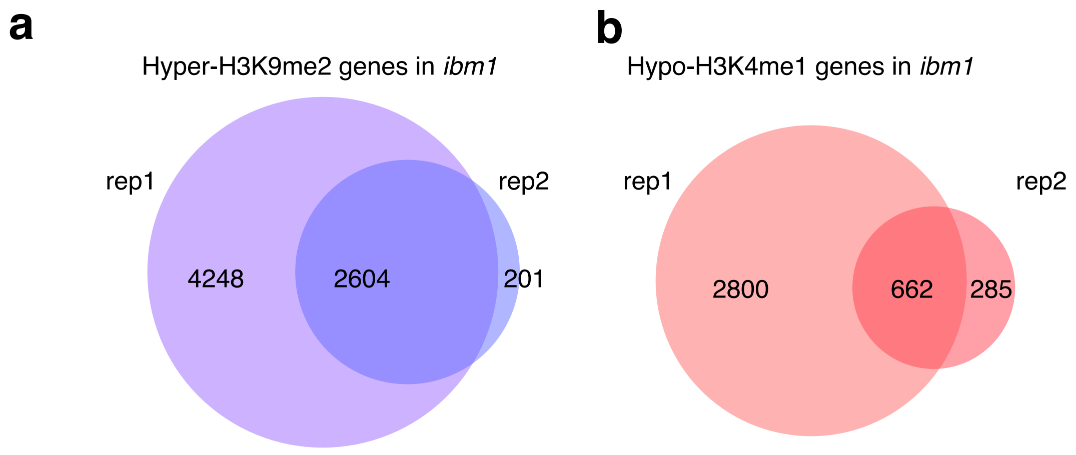

### Extended Data Fig.5

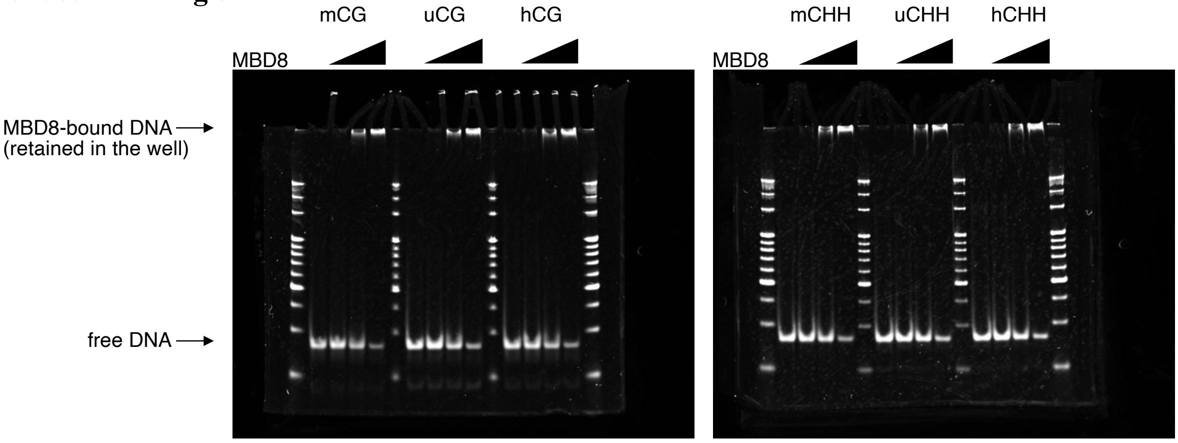


### Extended Data Fig.6

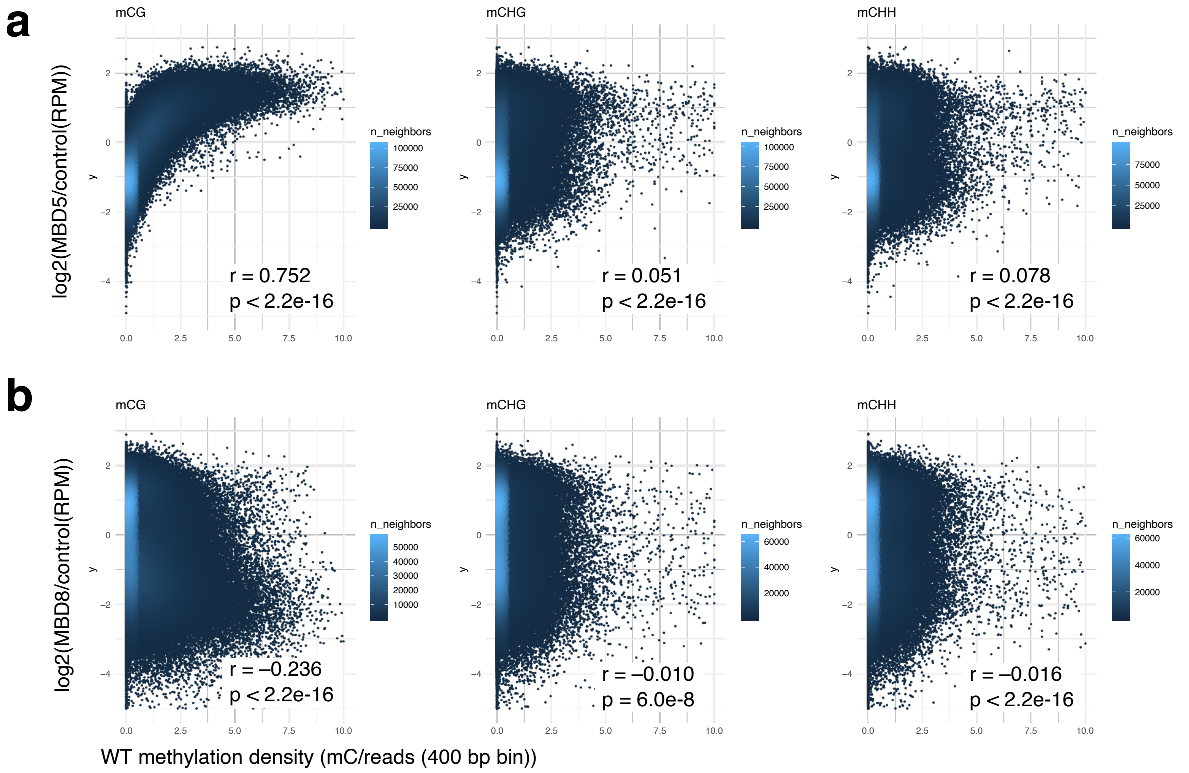

**Extended Data Fig.7**

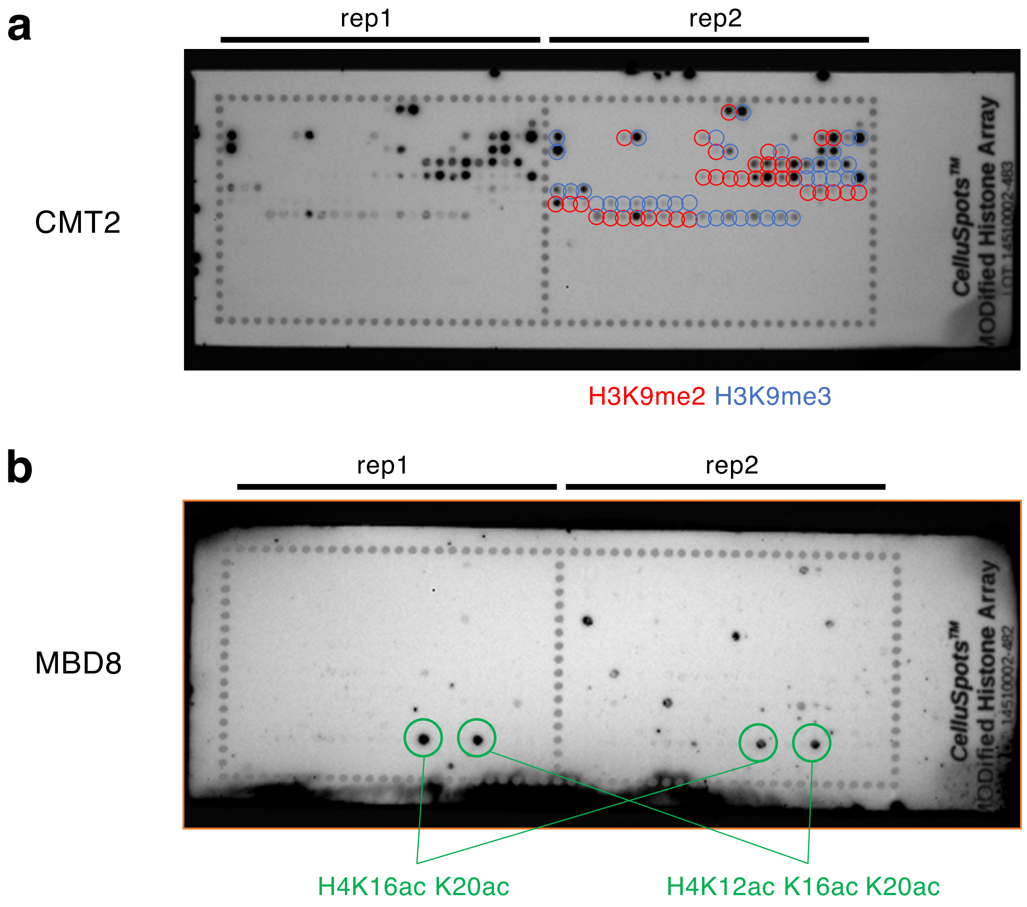

### Extended Data Fig.8

**a**

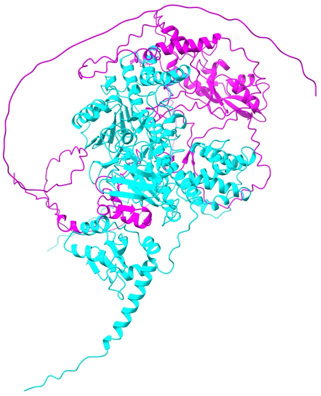

**b**

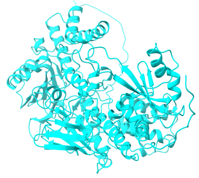

**c**

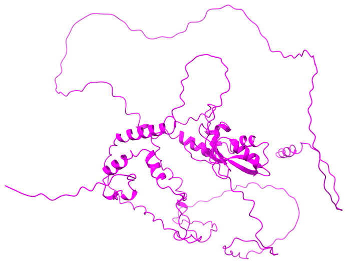

### Extended Data Fig.9

**a**

```
                                                  * * * *
LDL3 (875-934) SDVSYVSDVSAMDNSKHKVRVSTSNGCEYLGDAVLVTVPLGCLKAETIKFSPPLPDWKYA
 FLD (400-449) QTIRYGSNGVK----------VTAGNQVYEGDMVLCTVPLGVLKNGSIKFVPELPQRKLD
LDL1 (484-533) ESIRYGSNGVL----------VYTGNKEFHCDMALCTVPLGVLKKGSIEFYPELPHKKKE
LDL2 (375-424) DTIKYGDGGVE----------VISGSQIFQADMILCTVPLGVLKKRSIKFEPELPRRKQA
```

### Extended Data Fig.10

**a**

```
                                                        * * *
A. thaliana (1-50)          MDDGDLGNNHHNFLGGAGNRLSAESLPLIDTRLLSQSELRALSQCSSLSP
P. trichocarpa (1-46)       MATA----TVDSSICDLQNHLHIESLPLIDLRHLSQSELLSLSFCSSSPH
A. trichopoda (1-43)        -------MALPSDEKPDNPHLPISSIPLIDLRFLSQDEISSLALLSLPSS
Z. mays (1-30)              --------------------MGTEVAAVVDLRALTQSDLVALAAASPYAV
O. sativa Japonica (1-30)   --------------------MGTEVAPMVDMRALSQSDLVALAAGSPYSA
```
